# Gridlock from Diagnosis to Treatment of Multidrug Resistant Tuberculosis (MDR-TB) in Tanzania: Patients’ Perspectives from the Focus Group Discussion

**DOI:** 10.1101/402594

**Authors:** Stellah G Mpagama, Ezekiel Mangi, Peter M Mbelele, Anna M Chongolo, Gibson S Kibiki, Scott K Heysell

## Abstract

**Introduction:** Molecular diagnostics have revolutionized the diagnosis of multidrug resistant tuberculosis (MDR-TB). Yet in Tanzania we found delay in diagnosis with more than 70% of MDR-TB patients having history of several previous treatment courses for TB signaling complications of prior missed diagnosis. We aimed to explore patients’ viewpoints and experiences with personal and socio-behavioral obstacles from MDR-TB diagnosis to treatment in an attempt to understand these prior findings.

**Methods:** The study was conducted in December 2016 with MDR-TB patients admitted at Kibong’oto Infectious Diseases Hospital. We used semi-structured interviews and focus group discussion to examine patients’ views and experiences during MDR-TB diagnosis. Groups were sex aggregated to allow free interaction and to gauge gender specific issues in the social and behavioral contexts. The discussion – explored patients’ delivery factors that were impacting delay from MDR-TB diagnosis to treatment. Iterative data collection and analysis was applied with data, codes and categories being compared and refined.

**Results:** Forty-five MDR-TB patients participated in 6 focus group discussions. Challenges and barriers contributing to the delay from MDR-TB diagnosis to treatment were as follows: 1) The general population had differing understanding of MDR-TB that led to seeking services from traditional healers; 2) Also socio-economic adversity made health-seeking behavior difficult and often unproductive; 3) In the health system, challenges included inadequacy of MDR-TB diagnostic centers, lack of knowledge on behalf of health care providers to consider MDR-TB and order appropriate diagnostics; 4) Insufficiency in the specimen referral system for early diagnosis of MDR-TB. Non-adherence of TB patients to first-line anti-TB drugs prior to MDR-TB diagnosis given the multitude of barriers discussed was coupled with both intentional and unintentional non-adherence of health care providers to international standards of TB care.

**Conclusion:** Patient-centered strategies bridging communities and the health system are urgently required for optimum MDR-TB control in Tanzania.

## Introduction

Emergence of multidrug-resistant tuberculosis (MDR-TB) in settings with high burden of human immunodeficiency virus (HIV), diabetes mellitus (DM), and malnutrition poses an ominous challenge and threatens to dismantle the previous achievements in TB control [1]. MDR-TB is a laboratory diagnosis that requires not only identifications of *Mycobacterium tuberculosis* (MTB) but also determining resistance of at least rifampicin and isoniazid [2]. Therefore, diagnosis is either with culture and drug susceptibility testing (DST) requiring expensive biosafety level 3 or molecular diagnostics. Consequently, processes involved in the diagnosis of MDR-TB are complicated especially in resource-limited settings where laboratory services are not universally covered [3].

Furthermore, there remains a major gap between the numbers of patients diagnosed with MDR-TB compared to those stated on treatment. The World Health Organization (WHO) estimated in the year 2016, the global incidence of MDR-TB was 600,000. Yet in the same year, the MDR-TB global detection was 455,000 (78%) and only 205,000 (35%) enrolled for treatment [4]. In resource-limited settings where health systems are fragile, the gap is wider as illustrated in the WHO report of Tanzania for 2014. The report shows only 64 cases (10% of the estimated burden) of MDR-TB was detected and 28 (44%) was enrolled for treatment [5]. In Tanzania several signals have been highlighted such as a considerable delay in the MDR-TB diagnostic processes to treatment, and previous history of multiple episodes of TB retreatment; consequently resulted into few HIV and children treated for MDR-TB [6-8]. The low detection and treatment gap is the primary hindrance in the MDR-TB control in minimizing morbidity and mortality but also in preventing transmission in the community.

Although molecular diagnostics innovation such as XpertMTB/RIF and GenotypeMTBDRplus was considered as a major breakthrough for early MDR-TB diagnosis and treatment, its impact might have been diminished with other implementation factors [9]. We showed in Tanzania as others have elsewhere that roll-out molecular diagnostics had little difference in treatment outcomes compared with conventional methods despite a faster time to MDR-TB treatment initiation [8]. Therefore, we designed the current nationwide study to investigate the challenges and bottlenecks associated with delay in diagnosis to treatment of MDR-TB, both in patients -and health system, in a project supported by the World Health Organization. The expected research compendium branded as “gridlock from MDR-TB diagnosis to treatment” will provide evidence-based understanding of the magnitude of these delays in MDR-TB diagnosis, and its contribution on diseases transmission subsequently informing on the design of the interventions for breaking through such gridlock. In this second report from the compendium presented in this manuscript, we sought to understand these factors from patients’ experience. We specifically selected this qualitative research design to complement the quantitative investigations of HIV and TB clinics and diagnostic laboratories to better unravel the complexity of problem that is so depended upon individual and community interaction yet administered with nationwide or regional policies.

## METHODS

### Research design and Context

We used a focus group discussion (FGD) – a qualitative approach to understand patients’ experience during the disease period and processes they went through for seeking diagnosis to treatment of MDR-TB. A written informed consent (ICF) for FGD was provided to participants admitted at Kibong’oto Infectious Diseases Hospital (KIDH). Participants were invited to attend the FGDs if meeting the following inclusion criteria: Age of 18 years or more, Karnofsky score of 60 or more, treated MDR-TB for at least 2 months, and sputum smear negative. If participants could not read and write, the information sheets and consent forms were read out to them in presence of an impartial witness and a thumbprint was taken.

FGD was conducted at KIDH, which is a center of excellence for MDR-TB management in Tanzania. KIDH started MDR-TB treatment in 2009 and until 2016 it was the only center in Tanzania that managed MDR-TB cases. Recently the country has scaled MDR-TB services and new centers have started in 13 different regions. During the conduct of FGD however, MDR-TB referrals were from throughout the entire country and the average bed occupancy was 60 - 80 per day.

### Sampling strategy and data collection

Each FGD session consisted of 6-10 individuals. Groups were sex segregated to allow free interaction and to gauge gender specific issues in the social and behavioral contexts. On average most of the sessions lasted from one to one to two hours.

FGD with MDR-TB patients were held in a quiet hall located within hospital premises away from the patients’ wards. The hall had adequate ventilation (to minimize transmission) while retaining privacy. After initial introduction only research staff remained with participants. EM and SM facilitated FGD. EM is a sociologist with postgraduate training in qualitative research methods and has extensive experience in conducting FGD in health services in Tanzania while SM is a medical doctor with expertise in MDR-TB and clinical research. We took a systematic iterative approach to the development of FGD guide, which consisted around 5 open-ended questions. The discussion explored patients experiences encountered prior to diagnosis of MDR-TB, perceived contributing factors for delay in patients’ perspective. We also explored TB service delivery factors that were impacting delay in MDR-TB diagnosis and eventually amplification of the epidemic. We explored opinion from patients on ways and suggestions for improvement. We digital audio recorded the discussions and a dedicated note-taker took pertinent issues for follow up as they emerged during the discussion. The digital-recorded discussions and IDIs were transcribed verbatim in Kiswahili and thereafter translated into English.

### Data quality assurance and analysis

Transcription and audio recordings were reviewed for accuracy prior to further content analysis (by EM and SM) using the conventional approach of Hsieh & Shannon [10]. Clear patterns and discrepancies that emerged were discussed and consensus was reached among the study team, in many cases with review of the audio recordings. The main themes were presented to the study participants and further consensus was reached. This manuscript is reported according to the Consolidated criteria for reporting qualitative research (COREQ)

### Technical and Ethical Approval

Scientific and ethical approval was obtained from the World Health Organization Ethical Review Committee and the local Institutional Health Research Ethics Committee for author’s institute. For confidentiality audio-recorded tape and notebook was kept confidential and only authorized people accesses the materials. The tapes were destroyed after transcription into hard copies. This manuscript is reported according to the standards for reporting qualitative research [11].

## RESULTS

Sixty-two MDR-TB patients were admitted during the period of enrollment. Forty-five (73%) consented to participate and of those 5 did not participate due to medical severity. From the 40 total participants, six FGDs were arranged; 4 male and 2 female sessions were held in October 2016. Fortunately all participants in groups stayed for the whole session and no negative complication was reported even after the FDG. General clinical descriptors and demographics of the participants are summarized in table 1. Findings that contributed to delay from MDR-TB diagnosis to treatment were summarized and organized into 2 components: patients/population and health system factors. Incidental findings on the risk factors for developing MDR-TB are also summarized in the same manner

**Table 1.** Distribution of demographic and clinical characteristics of patients with MDR-TB participated in the focus
group discussion.

| Characteristics | Sub-categories | Number (%) |
| --- | --- | --- |
| HIV Status | Positive | 16 (40) |
|  | Negative | 24 (60) |
| History of previous TB treatment | None | 11 (28) |
|  | Treated TB once | 13 (32) |
|  | Treated TB twice | 10 (25) |
|  | Treated TB three times | 4 (10) |
|  | Treated TB four times | 2 (5) |
| Region of Domicile | Dar es Salaam | 11 (27) |
|  | Geita | 6 (15) |
|  | Mbeya | 5 (13) |
|  | Others category A | 8 (20) |
|  | Others category B | 10 (25) |

Category A: Mwanza, Mara, Njombe and Mtwara Regions each contributed 2 participants.

Category B: Morogoro, Kilimanjaro, Manyara, Shinyanga, Arusha, Singida, Sumbawanga, Tanga, Simiyu and Kigoma Regions each contributed 1 participant.

## 1. Factors in Delay from MDR-TB Diagnosis to Treatment

### 1.1. Patients/Community factors

The identified challenges and barriers contributing to delay from MDR-TB diagnosis to treatment initiation (Table 2) are further categorized into the following sub-themes:

**Table 2.** Summary of barriers and enablers for MDR TB diagnosis

| **Enablers** | **Barriers** |
| --- | --- |
| Social support | Most private facilities not testing TB |
| Isolation of MDR TB patients as advised by nurses | Failure to identify TB symptoms |
| Some patients are complying with treatment | Bread winner and single parenting- especially for female patients |
| Support from medical community | Traditional healers unaware of MDR-TB |
| Transport support to TB centers | Treatment without testing |
| Referral for further testing of even a suspected patient | Medical staff asking for money |
| Medical staffs making follow up for their patients to make sure that they go to test for TB | Private hospitals/health facilities have limited capability to diagnose TB, MDR-TB |
| Some nurses registering patients from far so that they can get treatment | TB clinics located close to HIV/AIDS offices |
| Provision of TB treatment at low cost | Non adherence to TB & MDR-TB treatment guidelines |
| Patients sacrificing other things for treatment | Providers unwilling to screen TB patients with persistent symptoms |
| Courage from patients to seek for TB health care | Lack of finance to send specimen for MDR screening |

#### 1.1.1. Knowledge Attitude and Practice

Findings suggest the community has limited understanding of TB in general and the additional gravity of MDR-TB. *“Majority of patients doesn’t have adequate knowledge thus they end up ignoring the situation” (FGD females*). Other patients do not easily accept the diagnosis of MDR-TB. For instance, a diagnosed MDR-TB patient secretly relocated, changed personal identifications, pretended to be a new case to another facility, was diagnosed with TB by smear microscopy but did not have additional drug-susceptibility testing performed and was thus started on first-line TB treatment. One participant narrated, *“I have used medications four times. Since I was still coughing, I decided to go to test at Musoma and results showed I had Chronic TB. I stayed for one week then I received a call that the bus has arrived and it took me to Moshi. I escaped after staying for two months to my mother who is staying at Simiyu. I was tested and diagnosed with Chronic TB. Thus they brought me back to Moshi for treatment” (FGD Males).* Other factors that facilitate patient delay include pharmaceutical shops as they provide drugs that relieve some symptoms as confessed, *“Honestly I don’t know regarding delaying as one may start coughing then ending up taking different medication, suspecting malaria. Until you start getting fever that’s when you go for testing and diagnosed with Chronic TB”* (FGD females). Alternatives to biomedical explanations of symptoms were common, even after MDR-TB diagnosis, and many patients sought traditional healers as they suspected they were bewitched. One patient revealed, *“After it reached a point that my condition became worse my relatives decided to take me back home. At home they took me to traditional healer but the moment I ingested the given medication I vomited it as it was” (FGD, males).* Other patient moved away to another village with concerns that he had been bewitched. *“We didn’t know I had TB, but I moved to another village thinking it was witchcraft. Since I was still coughing even after moving to another village for traditional medicine, then I opted to go to the hospital and tested sputum but I kept on believing it was witchcraft”* (FGD males).

#### 1.1.2. Social – economic adversity

Financial hardships affected the majority of participants and made their health seeking difficult and often fruitless. Several could not afford to pay for tests that ultimately resulted in delay in MDR-TB diagnosis and/or referral for treatment. For example a patient was not responding to first-line anti-TB treatment and was requested to have a chest X ray which was available only at another facility and was required to pay *“They told me to go get the treatment at a nearby health center, I really had to pay for the X-ray which costed me 16000/=TSHs (∼ 7.3 USD)” (*FGD males). One patient was unable to pay for transport cost after being referred for MDR-TB treatment (FDG male), *“ I completed 6 months of TB treatment though symptoms were persisting and I was referred to a regional hospital. I didn’t have money to cover for transport. I didn’t take my referral instead I went home to seek for support. After several days a friend of mine supported 10000 shillings (∼ 4.6USD) which supported me transport and I was diagnosed as MDR-TB.”* Some patients understood that costs they would incur would be catastrophic for their families, and while they understood their illness they tried to conceal it for as long as possible *“I am independent, I am paying rent myself, and I have a small kid and am working with Indians. This means not going for work, no job …” (FGD Females).* Another FGD female reported, *“ I am the day worker and I have twins younger than 7 years, I am the only one providing food and shelter, now I have MDR-TB I can’t support my family and I hesitated to start treatment”*

### 1.2. Health System

Although there were some examples of how the health system led to early diagnosis and referral, for example in tracing of contacts of previously diagnosed cases, *“While I was escorting my brother, then a nurse advised us to test for TB… I was also tested positive for MDR-TB thus I also started the medication immediately”* (FGD Females), the majority of participants reported struggles with the system and those findings were categorized into the following sub-themes.

#### 1.2.1. Inadequacy of MDR-TB diagnostic centers

For most patients when MDR-TB was clinically suspected, they were referred to distant facilities. *“They didn’t make me aware of my situation as it took me too long to know my condition”* (FGD Males). Others received different treatment for a long period of time prior to diagnosis of MDR-TB. *“I used to get recurrent fever, but at the private hospital where I was going they were testing me for malaria and UTI. As I started coughing and getting chest tightness the doctor advised me to go at the Regional hospital for sputum test…I was diagnosed with Chronic TB and I had to go to Kibong’oto for treatment”* (FGD Males). Other patients reported diagnostic equipment was non-functioning or unable to be used due to stock out of consumables.

#### 1.2.2. Lack of knowledge of health care providers to presume and test for MDR-TB

Patients on both first-line initial treatment and retreatment regimens presented often at the hospital complaining of persistence of TB symptoms despite of prolonged exposure to first - line anti-TB drugs as experienced in FGD female. *“I started getting fever, headache and I had severe chest pain too, I was tested and told I have Chronic TB then I was kept on treatment for six months without getting relief. Then I decide to go back to the hospital and I was given 60 injection. When I came back I was asked to collect sputum and they took the specimen to Mwanza*” (FGD females).

Further sentiments revealed an unwillingness of health care workers to test for MDR-TB and the use of improper inferences to defer diagnostic evaluation of patient who was not responding to first line anti-TB drugs. For example, HCWs briefly evaluated patients and said they look healthy and are fine without offering MDR-TB diagnostics, *“At the district the system is not good especially in testing of sputum. As one may go there for testing then they will start giving excuses that they don’t test sputum or they look at the general condition of a patient, if you don’t look sick, they will say that you are absolutely fine”* (FGD Males). Patients also perceived laboratory staff to be disinterested or giving excuses for not carrying out diagnostic procedures, *“You may find the lab technicians who test people for either Malaria, typhoid or HIV/AIDS you go get tested then they may tell you they are tired so most of them around here are young men”* (FGD Males). Others were more assertive HCW as they felt that the prescribed medication was not relieving them and they asked for another diagnosis and test. *“I confronted her again; she then said for now we have to wait for the Dr to come take you to Mbeya so that you may get tested again”* (FGD female). Other MDR-TB suspected cases were asked to pay for chest radiographs beyond the standard estimated fee for the government hospital *“I asked the doctor so, how can you help us? The only way I can help you is by first going to take an X-ray, of which you can come for the X-ray at around 4pm. So when we got there he said we should pay 100,000/= TSHs (∼ 44$) for the both of us (2 MDR-TB)”.* Another important factor was the tendency of other HCWs not to trust results from other health facilities and defer decisions to their own repeated testing which complicated the processes for the patient and often unnecessarily increased the cost to the patient and time to treatment. One patient argued, *“I told them I tested and results showed I have TB and I have come to take medicines. Then they told me they couldn’t put me on medication basing on the result of the place I was coming from, so I have to do another test then if the results will show I have TB then they will put me on medication. I agreed*” (FGD Males). Another participant showed clear symptoms of treatment failure after beginning first-line anti-TB drugs. The disease advanced from nodes to the lung, and the patient unfortunately received prolonged first line anti-TB treatment prior to MDR-TB diagnosis and referral as narrated in box 1.

#### 1.2.3. Inefficiency in the specimen referral system for early diagnosis of MDR-TB

Patients were required to deliver their own sputum to distant laboratories or were referred to a distant health facility close to a laboratory with the capacity of performing MDR-TB diagnosis. *“They told me that I have recurrent TB and I am supposed to produce sputum for DST and send myself to the diagnostic laboratory around 100 km. However I shouldn’t bother to follow the results because the laboratory will communicate the results to them. I travelled to that laboratory and left sputum after 3 days I was got a phone call from the hospital that I have MDR-TB ”*(FGD male). Also another patient submitted their own sputum to a laboratory approximately 50km laboratory with a capacity for MDR-TB diagnosis *“I was told to produce sputum for MDR-TB diagnosis while HCP were filling the laboratory form. I came back with specimen and I was told to take the specimen and the form. HCP directed me to travel to Tanga city to send the specimen and the form. I went outside to look for friends who could support me some money for fare. I took specimen and form and travelled to city …… the following week I needed to travel the same distance to trace for results. I took the laboratory results and brought to the facility and the HCP interpreted the results that I had MDR-TB” (FGD male).*

Other participants experienced multiple challenges in the community and within the health system, which covered several subthemes as highlighted in box 2.

## 2. Risk factors for developing MDR-TB

### 2.1. Patients/community

Shared behaviors in the village such as communal locations for alcohol consumption was postulated to contribute to TB spread, but also preventable death, *“Honestly speaking, back in the village people are so ignorant since you might find in the local bars people share the same alcohol containers. We blow and sip from the same containers… At the end of the day we keep on spreading the disease”* (FDG Males). It is almost impossible to prevent elderly from sharing the same container even with those who are sick. *“So when you find those old chronic drunkard men who drink the traditional alcohol out there in streets, it’s really hard to stop and prevent them from spreading the disease. There are so many people dying out there in the streets no joke I’m telling you”* (FDG Males).

### 2.2. Health System

Non-adherence to first-line drug-susceptible TB regimen was common and often due to behaviors for which the health system has little support (eg. mental health disorders, substance use), or a consequence of direct health system shortages (eg. medication stock-outs) *“I was drinking alcohol therefore there were times I stopped taking my medication so I went on like that for a while, there were weeks I was taking the medication and there were weeks I was not taking them at all”* (FGD Males). Furthermore, non-adherence behavior was brought by lack of social support or more episodic contact with the health system. “*Sometimes you forget to take them when you get family problems while you are at home hence you might stay a week without taking medication. Or otherwise you only remember to take them when you think of the nurse who gave you the medication or perhaps when you start coughing” (FGD Males).* Other non-adherence behavior was due to unrealistic expectations the health system has placed on patients, particularly in the context of medications administered by injection. *“… I just finished the dose without completing the injections as I was supposed to buy a syringe and I hadn’t monies. My condition got worse thus they took me to the Regional hospital where I was tested positive for MDR-TB” (FGD, males)*.

Direct lack of TB medications in nearby health centers resulted non-adherence for some patients. Patients explained that after testing positive for TB, patients were given medications for home based DOT. They were instructed that after finishing those medications, they were asked to refill in nearby health centers. One participant said, *“I was told if these medications finish, I should go and refill at a nearby health center within my village. After going there I was told that those particular medications are not available there”* (FGD male). Other health facilities had inconvenient service hours that prevented them from not getting the medications on time, *“Where I used to go and refill my medication, sometimes they tell me they are not opening on that particular day or perhaps they say today is Sunday so we don’t give out medicines while they are the one they gave me that date”* (FDG Males). Other facilities even denied the patients TB medications explaining that the patient was not assigned to that facility.

In some cases, health care providers tried to work outside the system to improve care, even to the point of falsifying the patient’s home location in the registry to assure access to anti-TB drugs. *“I had to go back to the nurse asking for her help. Then she said she is only helping me because I am sick, therefore she signed me up as a member of that region as a favor. … She filled the form and gave me the medicine and told me come back after two weeks to refill. Therefore I had been receiving the treatment for 6 months…”* (FGD male). In other scenarios, the absence of the health care workers from assigned posted, especially clinicians that in the current system are needed for authorizing diagnostics and approving refills, eventually results into non-adherence. *“So when I reached the hospital I never found the doctor, I went there twice but unsuccessful. Unfortunately I couldn’t go anymore because of the expenses. Then I had to go back to the nurse asking for her help”* (R7). Another participant added, *“They say even the doctors are not available or they may even say they are out off pills so they refer you … you find them closed. Unfortunately I ended up going back home without any medication. I was then admitted for Malaria treatment then I was lucky enough to be tested and confirmed positive for MDR-TB” (FDG Males).* Even after diagnostics are authorized and completed, it may require further authorization to release results to the patient. *“At the district hospital I was asked to give sputum for testing. I stayed for one week without been given results. They were saying doctors were not around, other days I found too many patients or already closed”* (FGD males).

*In addition to direct health system failures, evidence of blatant corruption was highlighted, ““… You may find him already drunk and he is saying that you should add more money at least 5000, then he gives you the medications and tells you to come back for refill and he insists that you should only go to him for the refill”* (FGD Males). Additional signal of corruption, the participant explained; *“Whenever you go back for the refill you find someone else and they start asking about your information all over again. They say they can’t see your file meanwhile your condition keeps on worsening as for some days they tell you they are out of pills* (FGD Males).

Stigma within the community was less commented on as expected but was referenced by on FGD male, *“ To blind the community who may directly link TB with HIV, majority of TB patients walk stealthily for anti-TB refill as you know TB clinics are very close to HIV clinics.”*

### Patients’ suggestions and recommendations for improvement

Participants highlighted general areas for improvement and specific mechanisms to decrease delay in MDR-TB diagnosis. Example increase screening and education in the community. *“In the village people are diagnosed very late so if possible they should pass house by house for testing” (FGD females*). Further suggestions were made for mass education in congregations, *“Apart from using billboards, education should be provided at areas with large population or mass e.g. at the market. Most of us get TB and other transmitted diseases there”* (FGD males).

The health system should increase diagnostic technologies and knowledge to health care workers. *“I think ivestigation should be done at the district and regional hospitals”* (FGD females).

## DISCUSSION

This study demonstrates how MDR-TB patients have been struggling tenaciously with their plight not only from the disease but also with the rigid and porous health system. Previously we quantified the delay in MDR-TB diagnosis from the time the specimen was submitted to treatment initiation [6-8], but in this detailed qualitative study patients themselves have identified noteworthy challenges that are rarely discussed by providers and policy makers but which severely limit any roll-out of new technologies or expansion of treatment programs.

Participants clearly raised concerns that the majority were now being treated for MDR-TB not because of the health system, but rather in spite of it. Frontline HCW routinely failed to suspect and test for MDR-TB suggesting that educational initiatives directed at HCW may have considerable programmatic benefit. Yet diagnostic delay was common even in centers that provided TB services, contrary to other countries where patients who attended TB centers had an advantage of being suspected and screened early [12]. Likely more important is that the health system lacks efficient mechanisms for onsite specimen testing or referral of specimens and relaying of results, a common phenomenon occuring in many developing countries [13], but one in which if ameliorated would concurrently raise awareness and testing for MDR-TB among the HCW. Lack of efficient mechanisms for specimen refferals was shown to contribute substantial delay in MDR-TB diagnosis to treatment similar to what other settings reports within and outside of sub-Saharan Africa [14, 15].

Importantly, participants described corruption as a major barrier and their accounts were vividly detailed inclduign health care providers asking for informal payment from patients for anti-TB refill and TB diagnostic tests. Some HCW indirectly gained financil benefit from sending patients for additional tests,such as x-rays at their affiliated diagnostics centers with five fold cost and without providing a receipt. The consequences included non adherence and delay in MDR-TB diagnosis to treatment as most of patients in the FGD suffered from severe economic hardship even prior to MDR-TB treatment. Corruption among those tasked for caring of patients with TB or other global health priority diseases compounds an already devastating diagnosis for an individual and on a larger scale has been likened to a biosecurity threat at the population level [16]. Our findings suggest that corruption among health care providers has contributed to the transmission of MDR-TB in Tanzania. Transforming the health system into a proactive state is urgently required not only for tackling TB but also other infectious diseases threats [17]. This will include community based diagnostic and treatment supppot, networking of health facilities at all levels through open access to diagnostics with transparent pricing, and strengthening public-private partnership to shed light on individual practices. A step toward this transformation may be to apply electronic continous medical education around priority issues such as TB and tie the medical education to licensing and recertification [18].

Contextualizing both health system barriers and patient/community level factors goes far to explain why most MDR-TB treated in Tanzania still occurs for patients that have had multiple courses of first line treamtent for TB. Our findings confirm that patients survive long and complicated courses from diagnosis to treatment, and represent a form of “long –term survivor” that likely represent only a fractional of the actual burden of MDR-TB in the community where many MDR-TB patients may have died before they access diagnosis or treatment [6].

Although we found effects on the mass media in driving the health seeking behavior in some of the MDR-TB patients who persistently sought for medical treatment [19], majority of patients and their family members did not know about MDR-TB presentation and correlated with alternative belief systems of illness such as witchcraft. Financial difficulty was also a barrier in many ways and included travel cost and living expenses during illness in particular for bread eaners. This had directly contribution in the delay of health seeking behavior mostly in those endured a prolonged illness [20]. Additional identified factors included inappropriate belief, stigma and dissatisfaction with the health system. In total participants describe a state of health seeking behavior for TB diagnosis that could be considerably improved but has changed little since the last national survey for such practices [21].

Patients’ perpectives of frontline helath care workers suggested substantial deficiencies in providers and clinics ability to implement essential l basics for directly observation therapy strategy (DOTS) that are fundamental for preventing acquired MDR-TB and transmission [22]. One of the elements in the DOTS requires TB treatment to be standardized with supervision and patient support [23]. Yet some patients confessed that they were non-adherent to the first line anti-TB drugs and practiced anti-TB drugs holiday for alcohol substitution. Others reported that they didn’t take anti-TB throughout because health facilities had no anti-TB medications, another additional DOTS element requiring an effective drug supply and management system [23]. Our patients’s perspectives expose just how difficult strict DOTS in TB endemic countries like Tanzania can be with unfunded or underfunded requirement, which at times were passed on to the patient in the form for extra cost or general hardship. In many instances our patient’s experiences were indirect conflict with the WHO guidance on the ethics of TB prevention, care and control which advocates for patients autonomy, diginity and integrity [24]. Indeed, patients’perspective highlight the need to rethink the necessity of DOTS as it is currently recommended and focus instead on community based solution to treatment [25]. Such community based and patients centered solutions are fitting with the current End TB Strategy [26] which calls for engagement of all relevant stakeholders, increase access to diagnostics, treatment and care for everyone with TB as a matter of not only public health but also social justice [27].

The study had limitations given that those patients that had successfully made it to diagnosis and treatment, therefore the responses may have been limited by survival bias. The ratio of FGD men and women was 2:1 suggesting ther may have been other factors driving a differential in MDR-TB diagnosis to treatment for women. One asumption might be men were prioritized by the family for treatment or diagnosed more at their place of work, for example limited TB screening efforts were underway in the mining industry. Nevertheless, one of the major strengths of the FGD was that participants were representative of the entire coutnry as at that time the facility was the only MDR-TB center for Tanzania. Furthermore the number of participants sampled of the total MDR – TB treatment population was relatively large. In addition we were able to identify some factors and practices that validated our previous observations.

## Conclusion

We have identified several contextual challenges and bottlenecks from our MDR-TB patients that cover both patient/community and health system factors along the pathway of MDR-TB diagnosis to treatment that contributed to substantial delay, and with both personal and public health impact. Current rollout of technologies like molecular diagnostics alone will not suffice in improving control of the MDR-TB epidemic, particularly in settings identified by this study’s participants. Urgent interventions directed at the patient-identified barriers could bring a greater return on TB care investment.

## Box. 1. FGD Participant describes the difficult pathway within the health system through MDR-TB diagnosis and ultimate treatment initiation

Unfortunately, initial suspicion of MDR-TB was very low. *“I had a diagnosis of TB adenitis and started anti-TB for 6 months. However at month 4, I developed cough and presented to the health facility where I was taking anti-TB. HCW responded that I should not worry; cough will subside because I am using medications”.*

Despite of the fact that cough and other constitutional symptoms were persisting the HCW did not consider MDR-TB as a cause of treatment failure. “*I presented to the same facility enquiring why cough was persisting. HCW emphasized that drug were working slowly and eventually symptoms will subside.*” When the patient completed anti-TB and after a week additional pulmonary TB symptoms manifested, there was still a lack of recognition by the health system of progressive TB disease. *“I experienced chest pain I thought to report to HCW however they told me that it is normal.”* The patient opted to visit a private health facility where he had chest radiograph and was informed that the picture showed active lung disease suggestive of TB. The private HCW advised the patient to report to the TB clinic and he opted to go to the same facility. He shared the report with the HCP at the same TB clinic and they opted to restart anti-TB without considering drug-susceptibility testing for MDR-TB. *“I started anti-TB and approximately after 2 months I met a doctor who asked me if I am using injectable agent, and I asked why?* Later, another HCW added streptomycin (an injectable aminoglycoside antibiotic) in to the failing regimen at approximately month 3. *“I completed 56 injectables and I was still coughing and experiencing chest pain, however I had other analgesics.”* Despite this, the patient did not have drug-susceptibility testing for MDR-TB but was instead referred to another facility for cardiovascular disease assessment including an electrocardiogram and was started on an anti-hypertensive. After completing 8 months of the second course of TB treatment there was no relief and thus presented to the same health facility, meanwhile cough and other constitutional symptoms were worsening. *“I told them I have well adhered to the treatment and I was told that symptoms will subside, I have stayed for 8 months and symptoms are persisting and I always use analgesics and anti-cough.”* After two months following the completion of the second course of TB treatment the patient experienced a dramatic weight loss of 12 kg. *“I was worried and the community was wondering what is happening to me. I opted to visit a private hospital and the private HCW advised me to visit.”* At that time pulmonary TB was again suspected and a sputum specimen was collected while the patient was given medication. After one week TB diagnosis was established by microscopy and the patient was recommended to begin first-line anti-TB treatment for the third time, but he refused while waiting MDR-TB testing. He was discharged to the community but within days he received a call and was informed that he has MDR-TB. The patient summarizes his struggle; *“Most likely I had MDR-TB since at the beginning of TB treatment. I feel HCWs are irresponsible and has lead me to suffer to this extent other wise I would have been treated this MDR-TB more than 15 months ago.”*

## Box 2. FGD participant experienced multiple community based and health systems challenges which contributed to the delay diagnosis of MDR-TB and treatment initiation

The patient sought TB services due to knowledge he gained through mass communication. *“I treated malaria at the private health facility (HF-1) however I felt I have TB because I am familiar with TB symptoms as I previously listened several times through the media.”* This resulted in presentation at the regional hospital (HF-2) where TB services are available. Although sputum smear was negative, diagnosis was made by chest radiograph. Appropriate first-line anti-TB treatment was started however for the first three months, symptoms were persisting. Patient reported to HF-2 and complained that symptoms were enduring. *“I have adhered treatment, not using alcohol, however the symptoms are not subsiding.”* HCP did not suspect MDR-TB and instead prescribed co-medication to treat symptoms, which bore additional direct costs to the patient. *“ I was supposed to pay 28,000 (∼ 12.2$) for medication however I was unable to buy drugs and I didn’t have money because I stopped working because of illness.”* The population/community is unaware of MDR-TB and alternative thinking prevails. *“My relatives were worried because I was taking anti-TB for more than 3 months without relief and cough was worsening, relatives compared to other known TB cases who had no symptoms within 2-3 months and were asking why me continuing with symptoms for this long. They though of witchcraft however I still believed that I had PTB. At the end of six month, I completed treatment without symptoms relief.”* The patient was understanding of the failure of the health system to provide further investigation, *“I never had monitored of sputum at month 2 or 3 or end of treatment. I am aware of the monthly monitoring.* Still *at the end of treatment I reported to the HCP and complained on the persisting symptoms. The HCP told me that they are done, if I feel I am still sick I should consult a doctor therefore I should restart the consultation process through the general clinic.”* In addition to the lack of diagnostics during and at the end of treatment, other opportunities were missed, “*I opted to go home however I felt slightly well and I resumed work however during the night I had excessive cough, nigh sweat and chest pain. I thought to go back at the same hospital but I lost trust. My wife told me that this is not normal TB most likely is related to witchcraft as anti-TB has not helped. Regardless I believed, it was TB I thought to travel to another district to my uncle to seek for support as he is slightly well off because I had financial difficult and I thought my uncle may send me to Muhimbili National Hospital. The uncle advised to visit a near health facility (HF-3) where I had a PTB diagnosis and HCP told me I had extensive lung damage. HCP recommended that I should submit sputum for MDR-TB diagnosis. I wondered I didn’t know about MDR-TB. However they told me I should send the specimen myself to Central TB Reference Laboratory because the dates for them to submit specimen have already gone. I took the sputum specimen and submitted to CTRL thereafter I started same medication I used before and injectable. Yet the symptoms did not subside and I asked for results they said not yet.”* Despite prior first-line TB treatment and extensive lung disease, the patient’s perception of the HCW was that they believed the urgency was low to test for MDR-TB, even directing the patient submit his own specimen to the Central TB Reference Laboratory. In actuality, the specimen was also not set up for rapid MDR-TB testing, such as with the Xpert MTB/RIF assay, but rather the conventional culture based methodology which takes months or more. The patient lost trust, *“I opted to run away from the health facility and shifted to another health facility (HF-4) where I needed to travel at least change 2 buses. The new health facility asked me to produce sputum and for about 5 days I went there for daily injectables but I got challenges in the bus as I was tired sometimes I don’t get a sit and I asked passengers to help me. I reconsidered that it is expensive to travel every day and I should surrender myself at HF-3. Fortunately at HF-4 work out sputum results and I was told I had MDR-TB diagnosed by molecular diagnostics and I was referred for treatment.”* This example highlights that prioritizing screening with molecular diagnostics could prevent morbidity and mortality related to MDR-TB, and also prevent the transmission of MDR-TB in high-risk settings such as long rides on public transportation. “*While I was on MDR-TB treatment for 4 months I received a call from HF2 that culture and DST results show that I have MDR-TB and results came out 8 months since I submitted sputum.”*

HF-1, HF-2, HF-3, HF-4 labels different four health facilities with HF-1 being the first facility visited followed by second, third and the last one respectively

